# Telomere Length in Plants Estimated with Long Read Sequencing

**DOI:** 10.1101/2024.03.27.586973

**Authors:** Kelly Colt, Semar Petrus, Bradley W. Abramson, Allen Mamerto, Nolan T. Hartwick, Todd P. Michael

## Abstract

Telomeres play an important role in chromosome stability and their length is thought to be related to an organism’s lifestyle and lifespan. Telomere length is variable across plant species and between cultivars of the same species, possibly conferring adaptive advantage. However, it is not known whether telomere length is related to lifestyle or life span across a diverse array of plant species due to the lack of information on telomere length in plants. Here we leverage genomes assembled with long read sequencing data to estimate telomere length by chromosome. We find that long read assemblies based on Oxford Nanopore Technologies (ONT) accurately predict telomere length in the two model plant species *Arabidopsis thaliana* and *Oryza sativa* matching lab-based length estimates. We then estimate telomere length across an array of plant species with different lifestyles and lifespans and find that in general gymnosperms have shorter telomeres compared to eudicots and monocots. Crop species frequently have longer telomeres than their wild relatives, and species that have been maintained clonally such as hemp have long telomeres possibly reflecting that this lifestyle requires long term chromosomal stability.

## Introduction

Telomere repeat arrays of a primary seven base pair sequence are essential components of all eukaryotic chromosomes. They protect the ends of chromosomes from exonuclease degradation and chromosomal fusions as well as eliciting DNA damage response (Procházková Schrumpfová, Fojtová and Fajkus, 2019; Peska and Garcia, 2020). In some species, telomeres are also implicated in the polar organization of chromosomes in the nucleus during meiosis known as the Rabl formation (Cowan, Carlton and Cande, 2001) and they may also play a role in the evolution of chromosomes through degradation and speciation fusions (Stindl, 2014). Therefore, the maintenance of species-specific telomere array length plays a role in not only maintaining the integrity of chromosomes but also that of the species. Vertebrates’ initial chromosome length is not strongly correlated with longevity, however longevity is strongly linked with average annual telomere shortening; the fewer base pairs lost annually, the longer lived the organism (Whittemore *et al*., 2019). This suggests that the robustness of a species’ DNA repair mechanisms are a determining factor in longevity (Whittemore *et al*., 2019). Larger species (>5kg) tend to have shorter telomeres and repress telomerase at maturity, which requires active mitigation of oxidative damage to preserve telomere length; Smaller species (<2kg), with exceptions, have convergently evolved longer telomeres and active telomerase, which costs less than oxidative protection mechanisms (Gomes *et al*., 2011; Risques and Promislow, 2018).

The relationship between telomere length and longevity among plants remains unclear; published minimum, average, and maximum telomere length can be highly variable between species, accession, chromosome, tissue type, and tissue age (Figure 1; Supplemental Table S1). Bristlecone pine (*Pinus longaeva*) is thought to be the oldest living non-clonal eukaryotic organism with the oldest tree on record named “Methuselah” being 4,765 years old. The average telomere length was 6 kilobases (kb) across several tissues (needle and root) and ages (37 to 3,500 years), while the max length varied substantially with the longest telomeres (32 kb) found in the roots of the 3,500 year old tree (Flanary and Kletetschka, 2006). Another long-lived gymnosperm that lives beyond 3,500 years old (y/o), *Ginkgo biloba* has an average telomere length of 3-4 kb (Shan-An, Gu and Zi-Jie, 1997; Liu *et al*., 2007). Long-lived (2000-5000 y/o) trees such as Coastal Redwood (*Sequoia sempervirens*), Rocky Mountain Bristlecone Pine (*Pinus aristata*) had longer average (15 kb) and max (50 kb) length telomeres vs medium-lived (400-500 y/o) Western White Pine (*Pinus monticola*) average (8 kb) and max (15 kb) (Flanary and Kletetschka, 2006). These results are consistent with other conifers tested, such as Norway spruce that has an average length of 9-23 kb (Aronen, Virta and Varis, 2021) as well as Scots Pine that has an average length of 15-23 kb (Aronen and Ryynänen, 2012). In shorter lived model plants and crops *Arabidopsis thaliana*, *Oryza sativa* (rice), and *Zea mays* (maize) telomere lengths are between 2-5 kb, 5-20 kb, and 6-25 kb respectively. In addition, there is variability among telomere lengths across different accessions within a species, while length is conserved within an accession (Shakirov and Shippen, 2004; Brown *et al*., 2011; Fulcher *et al*., 2015; Choi *et al*., 2021).

**Figure 1.**
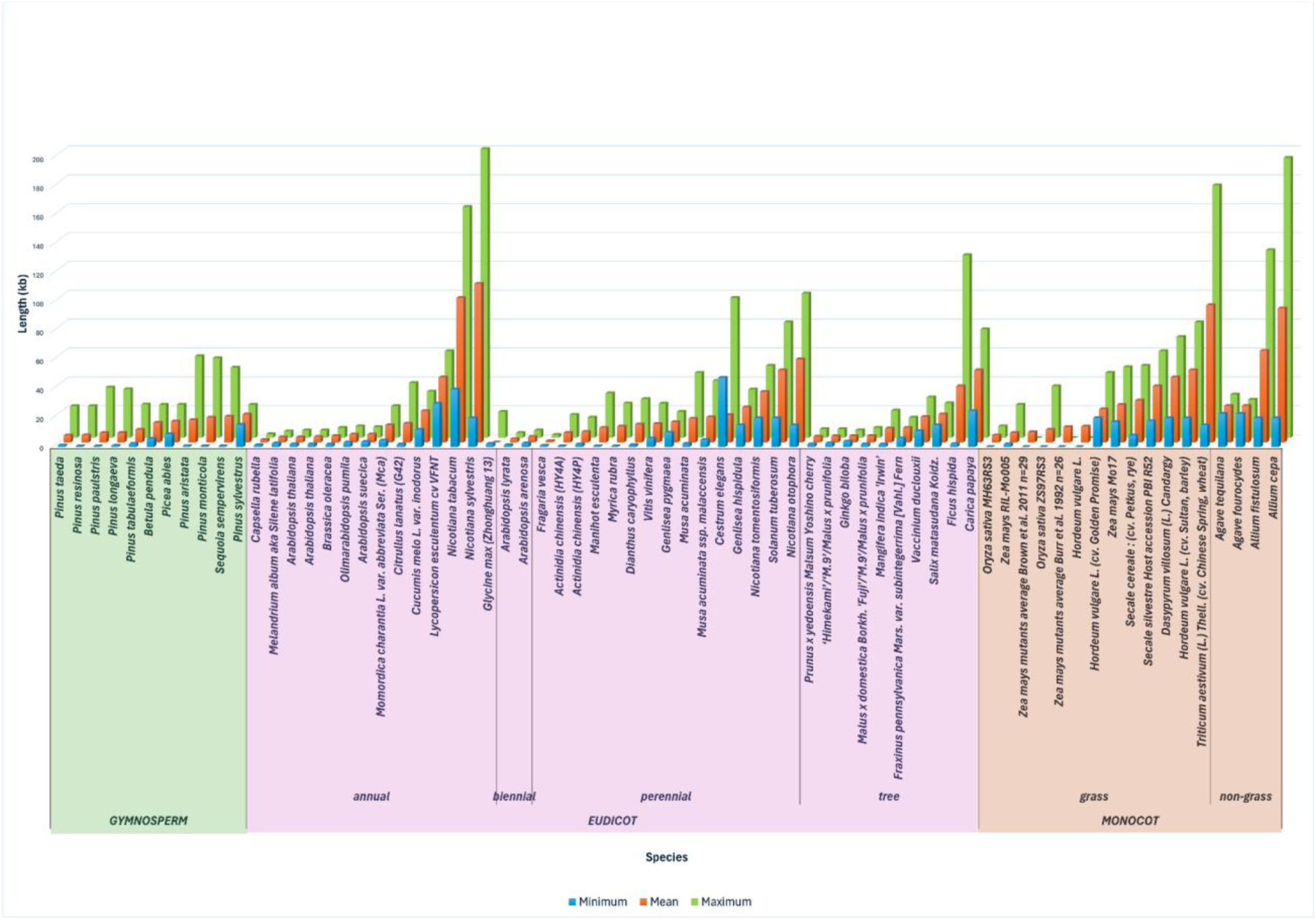
Published telomere length estimates in plants. Separated between Dicots and Monocots, Averages were taken for the *Zea mays* mutants from (Brown *et al*., 2011) and (Burr *et al*., 1992) for visualization. All values and additional data in Supplemental Table S1.

Telomere length in plants has been associated with life history traits including seasonal and phenotypic variation, environmental interactions, sex, flowering time, and seed quantity (Liu *et al*., 2007; Mu *et al*., 2014; Choi *et al*., 2021; Osnato, 2021; Campitelli *et al*., 2022). Recent work in arabidopsis showed a number of associations between the telomere length and environmental response, with effects on phenotype (number of stomata, osmotic potential, leaf hydration), seed production, and overall reproductive fitness; phenotype variations were directly proportional to the telomere length divergence from the wild type Col-0 (Campitelli *et al*., 2022). Recently it has been shown that telomere length is not affected when telomerase is increased either due to zero gravity (space flight) or artificially (Barcenilla *et al*., 2023). However, the role of telomere length and what controls telomere length is not known in plants.

Historically telomere length has been measured using a variety of methods, the most common being Terminal Restriction Fragment (TRF) analysis (Fajkus, Sýkorová and Leitch, 2005) (Supplemental Table S1). The assay involves digestion of genomic DNA using BAL-31 and other restriction enzymes, leaving telomere regions free for visualization by southern blot using fluorescent probes (Fojtová *et al*., 2015). *Arabidopsis* telomeres were originally measured using TRF analysis, indicating the telomere length was approximately 2-5 kb (Richards and Ausubel, 1988). Subsequently, the Cherry Tomato (*Lycopersicon esculentum* cv VFNT) and Tobacco (*Nicotiana tobaccoum*) telomeres were measured using TRF with resulting telomere lengths between 30-60 kb and 60-160 kb, respectively (Ganal, Lapitan and Tanksley, 1991; Fajkus *et al*., 1995). Rice (*Oryza*) telomere length and placement has been extensively studied using fiber-Fluorescent In-Situ Hybridization (fiber-FISH) and TRF analysis, revealing telomere lengths between 5-20 kb depending on the species (Mizuno *et al*., 2006).

More recently, quantitative PCR (qPCR) emerged as a more high throughput method to determine telomere length (Cawthon, 2009; Vaquero-Sedas and Vega-Palas, 2014). Despite these advances, the variety of telomere sequences complicates existing measurement techniques which are reliant on specific recognition sites for probes and primers (Weiss and Scherthan, 2002; Sykorova *et al*., 2003; Peška *et al*., 2015; Kumawat and Choi, 2023). As of 2017, telomere sequence is only known for approximately 10% of the plant species in the Plant rDNA database (Vitales *et al*., 2017; Peska and Garcia, 2020). Modifications in the telomere sequence are relatively common, frequently increasing the length and complexity of the repeated sequence (Weiss and Scherthan, 2002; Sykorova *et al*., 2003; Peska, Sýkorová and Fajkus, 2008; Peška *et al*., 2015; Tran *et al*., 2015; Fajkus *et al*., 2016; Peska and Garcia, 2020). Additionally, TRF analyses are hard to compare across studies given the variety of restriction enzymes chosen, and results are highly dependent on DNA and blot quality (Aubert, Hills and Lansdorp, 2012; Peška *et al*., 2015). FISH requires actively dividing cells, limiting the viable tissues in plants to assay, and qPCR results are notoriously difficult to reproduce (Weiss and Scherthan, 2002; Aubert, Hills and Lansdorp, 2012). Methods to more extensively, accurately, and reproducibly measure telomere length have not been developed until more recently.

Illumina short read whole genome sequencing is emerging as an alternative approach to establish telomere length (Paez, Holliday and Hamilton, 2023), and as long read sequencing technology continues to improve, telomere to telomere (T2T) genome assemblies have also become possible (see discussion). The advent of Oxford Nanopore Technology (ONT) long read single molecule sequencing has enabled the ability to resolve the length of telomere repeats directly in our genome assemblies. Strand specific errors in ONT base calling of telomere sequence have been resolved by re-training the Bonito basecaller with telomere specific data, resulting in highly improved accuracy (Tan *et al*., 2022). We have documented that the telomeres of the basal non-monocot aquatic Duckweed *Spirodela polyrhiza* have an average telomere length similar to that of *A. thaliana* at ∼5 kb (Hoang *et al*., 2018; Michael *et al*., 2018; Harkess *et al*., 2020). Therefore, we were interested to see if we could leverage ONT length read assemblies to look broadly across plants with different life-styles. In this study we apply our informatic pipeline to an array of plant genomes to estimate their telomere length. We then confirm these telomere length estimates with a new bioinformatic tool that enables direct estimation from reads as well as by qPCR. The work we present here sets that stage for more studies to understand how chromosome evolution and plant habit is influenced by telomere length and *vice versa*.

## Results

### Published telomere lengths across plants

We compiled as many published telomere lengths found in the literature as possible in addition to performing our own analysis (Figure 1; Supplemental Table S1). We noticed a potential trend that highly cultivated crop species including *Triticum aestivum* (wheat), *Nicotiana tabacum* (tobacco), *Cannabis sativa* (hemp), and *Allium cepa* (onion) have exceptionally long telomeres, while other crop species *Oryza sativa* (rice) and some *Zea mays mutants* (maize) are on the opposite side of the spectrum with relatively short telomeres. Cultivated wheat, tobacco and maize generally have longer telomeres than their wild relatives (Soreng: and Peterson, no date; Burr *et al*., 1992; Fajkus *et al*., 1995; Vershinin and Heslop-Harrison, 1998; Brown *et al*., 2011). Conversely, cultivated rye appears to have slightly shorter telomeres than its wild counterpart (Vershinin and Heslop-Harrison, 1998).

### Telomere length in long read Arabidopis thaliana

Several reports have shown that the *Arabidopsis* Col-0 telomere ranges from 2-5 kb (Shakirov and Shippen, 2004), and a recent study using a combination of K-mer repeat copy number estimation and the lab-based terminal restriction fragment (TRF) method showed that the average length across *Arabidopsis* accessions is 3,533 bp (Supplemental Table S1) (Abdulkina *et al*., 2019; Choi *et al*., 2021). We sequenced the *Arabidopsis* Col-0 accession with Oxford Nanopore Technologies (ONT) to finish the centromeres and this dataset was also useful to validate telomere length from a long read assembly (Naish *et al*., 2021). The resulting ONT reads had a raw N50 length of 22 kb (Figure 2A) and were scaffolded against TAIR10 using ragtag (Alonge *et al*., 2022). We identified two different types of centromere arrays in the Col-0 assembly with base units of 159 and 178 bp (Figure 2B); it has been shown in *Arabidopsis* that the 178 bp array is the active centromere (Maheshwari *et al*., 2017; Naish *et al*., 2021). We then searched the new Col-0 assembly for tandem repeats (TR) and we identified 18 telomere (AAACCCT or AGGGTTT) arrays with an average length of 2,734 bp, and the longest being 18,522 bp. Five telomere arrays were found on the ends of chromosomes with an average length of 2,760 bp (Figure 2C). The remaining telomere arrays were found proximal to the centromeres (10) or on unplaced contigs (3). These results were consistent with the other measures that the *Arabidopsis* Col-0 telomeres were between 2-5 kb, and suggested that the informatically derived telomere estimates could be skewed by internal telomere arrays since they made up 50% of the placed telomere arrays.

**Figure 2.**
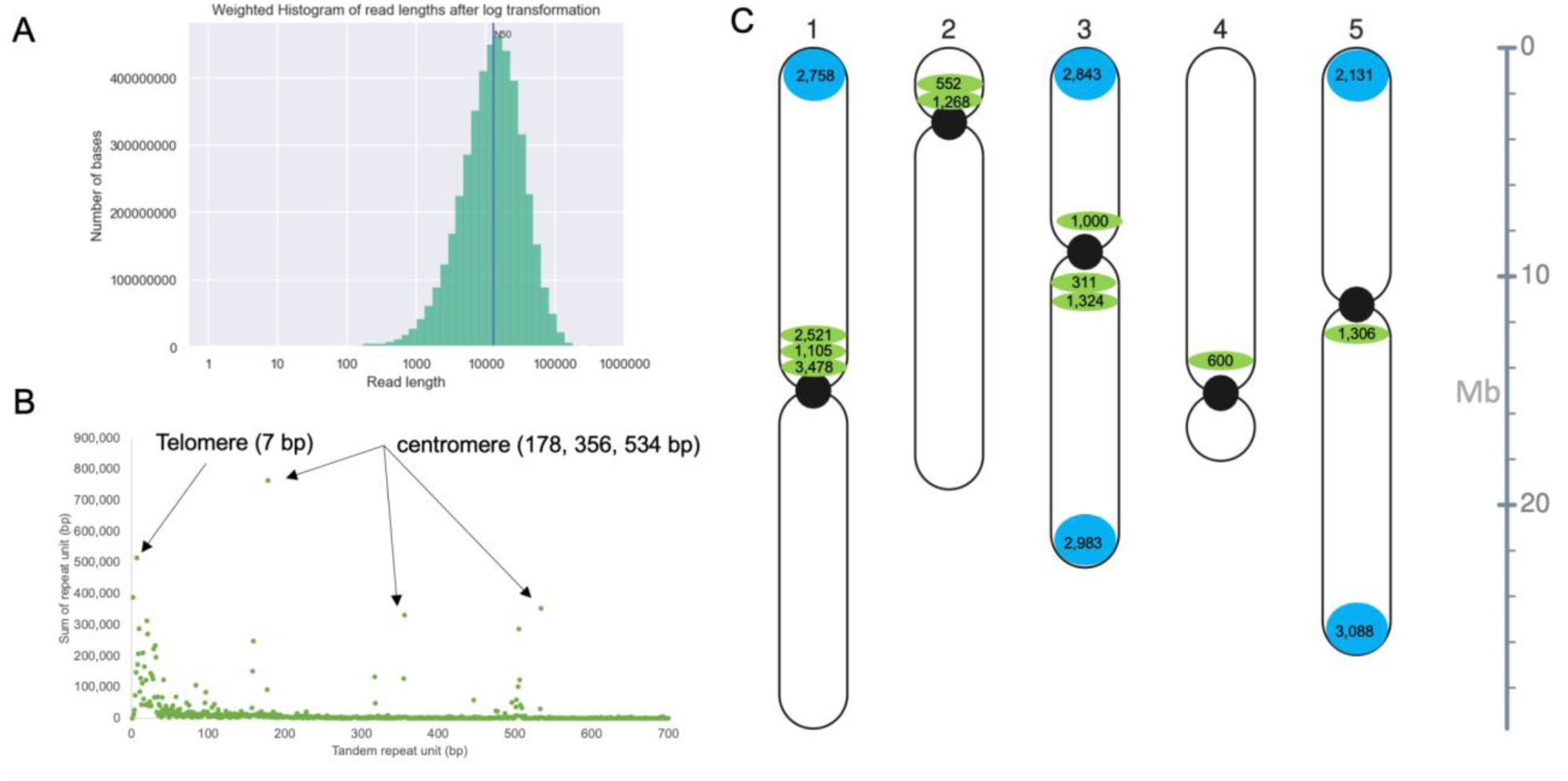
Long reads and long read assemblies enable the estimation of telomere length by chromosome in *Arabidopsis*. A) Weighted log transform histogram of ONT R10 read lengths with the read N50 length (12 kb) indicated. B) Telomere and centromere arrays identified using tandem repeat finder (TRF) and plotted by summed length. The telomere is part of the 7 bp repeat cluster, while the active centromere is at 178 bp with higher order repeats (HOR) at 356 and 534 bp. C) Arabidopsis chromosomes with the 178 bp centromeres (black circles), terminal telomeres (blue circles; >80% identity) and internal telomere sequence (green circles; <80% identity).

### Rice telomere length estimated in long read genome assembly

Rice telomeres are also well characterized (Mizuno *et al*., 2006; Mizuno, Matsumoto and Wu, 2018; Song *et al*., 2021), providing a second proof point as to whether estimating telomere length with long reads is an effective method. Rice telomeres are on average 5-10 kb but can vary from 5-20 kb across different varieties (Mizuno *et al*., 2006; Mizuno, Matsumoto and Wu, 2018). Two chromosome-resolved circum-basmati rice genomes were generated using ONT long reads (Choi *et al*., 2020), which we leveraged to estimate the length of telomeres across the rice chromosomes. The read N50 of the circum-basmati rice reads were ∼33 kb, which was longer than the Col-0 input reads, and the resulting assemblies were contiguous with a contig N50 length of 6.32 and 10.53 Mb (Choi *et al*., 2020).

Both assemblies had the canonical centromere array called CentO, which had a 155 bp base repeat unit and the non-active 165 bp array that was predominantly found on Chr09 (Figure 3A). The telomere sequence had an average length of 13,014 and 11,310 bp for Basmati344 and Dom Safid respectively, and 8 of the 12 chromosomes had telomere sequence on both ends of the chromosomes (Figure 3B). The longest array in basmati334 was 18 kb and most of the telomeres were properly placed at the end of the chromosomes, but several were located proximal to the end of the chromosomes (Figure 3C). This could be due to the method that the contigs were scaffolded with RAGOO (Alonge *et al*., 2019), or a reflection that only the canonical telomere sequence were searched for and early studies have suggested that other related arrays have been found in rice (Mizuno *et al*., 2006; Mizuno, Matsumoto and Wu, 2018). These results were consistent with previous measurements and suggested that estimating telomere length from long read assemblies was generally an accurate proxy comparable to other methods.

**Figure 3.**
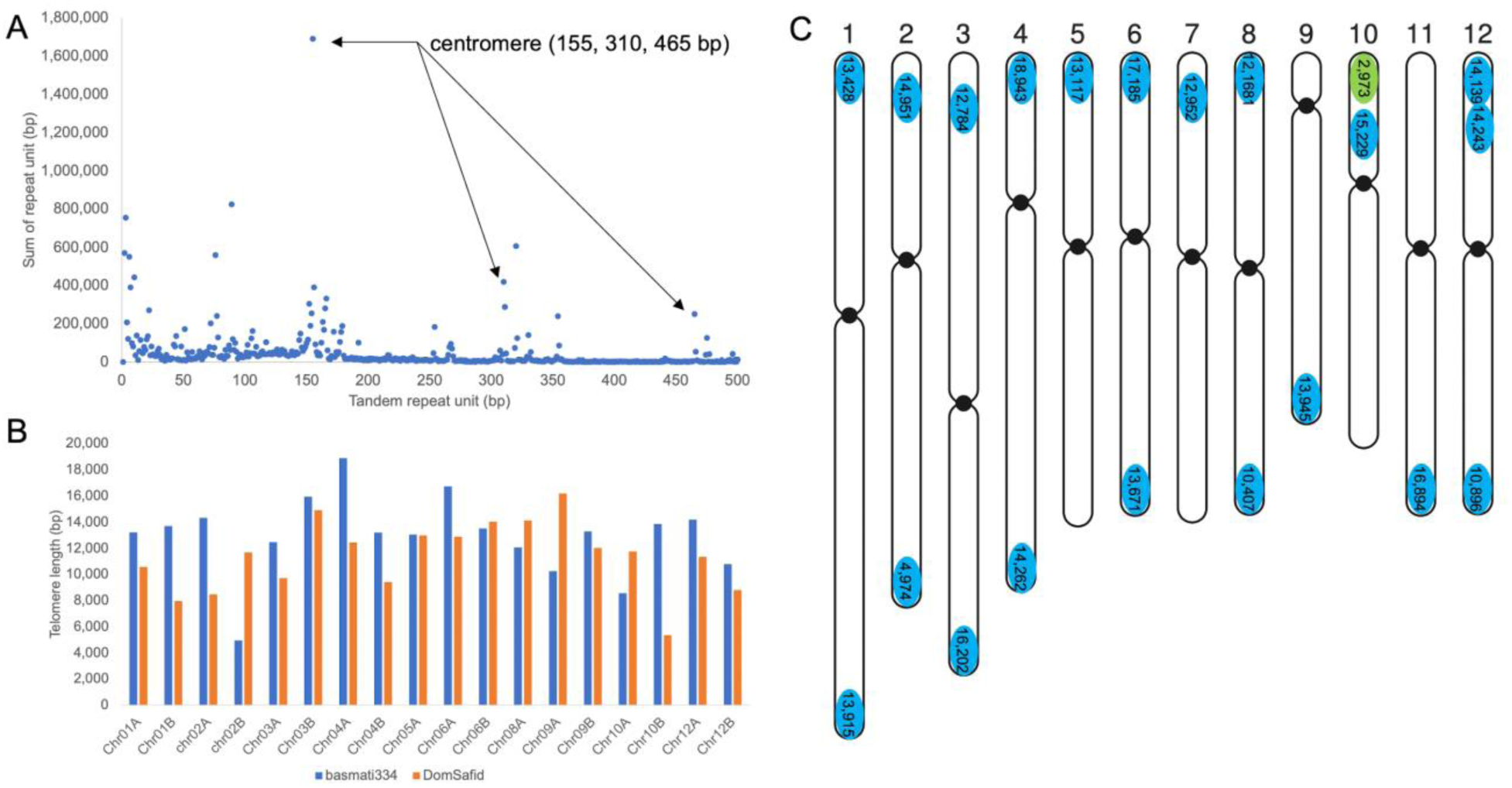
Long read assemblies enable the estimation of telomere length by chromosome in *Oryza sativa* (rice). A) Telomere and centromere arrays identified using tandem repeat finder (TRF) in the basmati334 genome assembly and plotted by summed length. The telomere is part of the 7 bp repeat cluster, while the active centromere is at 155 bp with higher order repeats (HOR) at 310 and 465 bp. B) Comparison of telomere length between the basmati334 (blue) and Dom Sufid (orange) genome assemblies. C) Basmati334 chromosomes with the 155 bp centromeres (black circles), terminal telomeres (blue circles; >80% identity) and internal telomere sequence (green circles; <80% identity).

### Estimating telomere length across ONT-based plant genome assemblies

Next we leveraged the long read assembly telomere estimation method to ask whether there is a relationship with plant lifestyle or lifespan. We identified ONT-based genomes from annuals that are close dicot relatives of *Arabidopsis* such as pennycress (*Thlaspi arvense*) and black mustard (*Brassica nigra*). We also identified other dicots such as the perennial woody crop cranberry (*Vaccinium macrocarpon*) and the annual, but predominantly clonally propagated crop, hemp (*Cannabis sativa*). For long lived dicots we identified several long-lived trees such as *Eucalyptus pauciflora* (Wang *et al*., 2020), Inga (*Inga sapindoides*) and the extremely long lived baobab (*Adansonia digitata*). In addition to the circum-basmati rice genomes, we identified ONT-based genomes for other grasses such as *Sorghum bicolor* (Deschamps *et al*., 2018) and fonio (*Digitaria exilis*). For non-grass monocots we found the annual herb banana (*Musa schizocarpa*) (Belser *et al*., 2018) and clonal duckweed species *Spirodela polyrhiza* (Harkess *et al*., 2021)*, Landoltia punctata (Baggs et al., 2022), Lemna minuta* (Abramson *et al*., 2021)*, Wolffia australiana* (Michael *et al*., 2020)*, and Wolffiella neotropica*. Finally, we also tested one of the longest lived gymnosperms, which only makes two leaves and lives for centuries, Welwitschia (*Welwitschia mirabilis*) (Wan *et al*., 2021).

Since we found that most telomere arrays at the end of the chromosomes in *Arabidopsis* and rice had high identity, we limited the comparison of telomere lengths across species to telomeres identified on the ends of chromosomes/contigs/scaffolds with an identify greater than 80%. Similar to *Arabidopsis* and rice, in banana we found typical centromere arrays with a base unit of 220 bp and HOR, as well as telomere repeats slightly larger than those of the grasses (Supplemental Figure S2). When we looked across the set plants several trends stood out (Figure 4). The annuals (rice and the brassicas) in general had the shortest average telomeres, and the long lived trees (inga and baobab) had long average telomeres, which is consistent with the idea that telomere length reflects longevity, even in plants. However, three unexpected relationships emerged. Hemp had the longest telomeres of all plants tested, the fast-growing clonal duckweeds displayed wide variation in telomere length and the long lived Welwitschia had telomeres similar in length to *Arabidopsis* (annuals).

**Figure 4.**
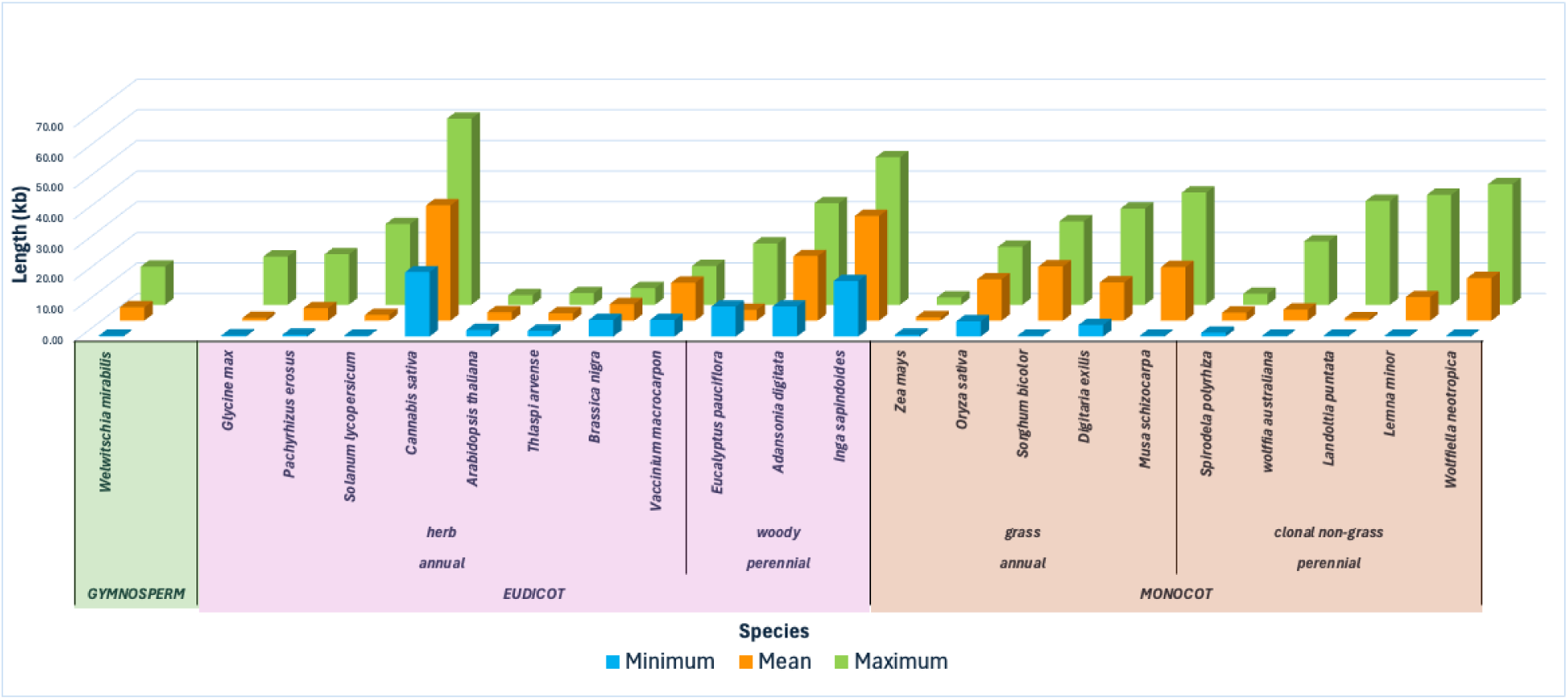
Estimated telomere length across plant species with different lifestyles and lifespans. The species are organized: dicot, gymnosperm, and monocot, and the average telomere length (green), shortest telomere length (blue) and the longest telomere length (green) are displayed for each species. Data is sorted by maximum length. The values and additional data can be found in Supplemental Table S2.

Several methods have been demonstrated to measure telomere length in the lab such as the terminal restriction fragment analysis (Abdulkina *et al*., 2019; Choi *et al*., 2021) and a quantitative PCR (qPCR) based method that enables rapid relative telomere length estimates (Cawthon, 2009; Vaquero-Sedas and Vega-Palas, 2014). Due to the cost and speed of the qPCR assay, and the power to use this assay within a specific plant to find relative changes in telomere length, we used this assay to validate the telomere size in *Arabidopsis*, baobab, *Welwitschia* and *Wolfiella*. The results are consistent with published results in *Arabidopsis* and the relative difference in telomere length that we found with both the genome assembly and raw read informatic analysis (Supplemental Figure S3). Together these results support the use of these different approaches to estimate telomere length across a diverse array of plants, which will lead to a better understanding of the importance of telomere length in plants.

### Estimating telomere length directly from raw ONT reads

Sometimes only raw long reads are available, either deposited in the short read archive (SRA) without a genome assembly, or there is a desire to generate skim sequences and estimate telomere length directly from raw reads. In the later case, skim sequencing would enable more broad sampling of the same plant for changes in telomere length. Therefore, we developed a pipeline for estimating telomere length directly from raw long reads called TeloNum (https://gitlab.com/semarpetrus/telonum). We estimated telomere length from the same raw ONT reads that we used to make the assemblies (Figure 5; Supplemental Table S2) to compare these two approaches. We found that the median and maximum estimated telomere length was correlated (0.56 and 0.84, respectively) (Figure 5A,B). The correlation was less strong for the median due to the variability in short estimates from the raw sequence, possibly due to truncations or reads well below the N50 of the dataset that would be extended during assembly. Therefore, in general we leveraged the maximum telomere length from TeloNum as a gauge of the telomere potential in a plant.

**Figure 5.**
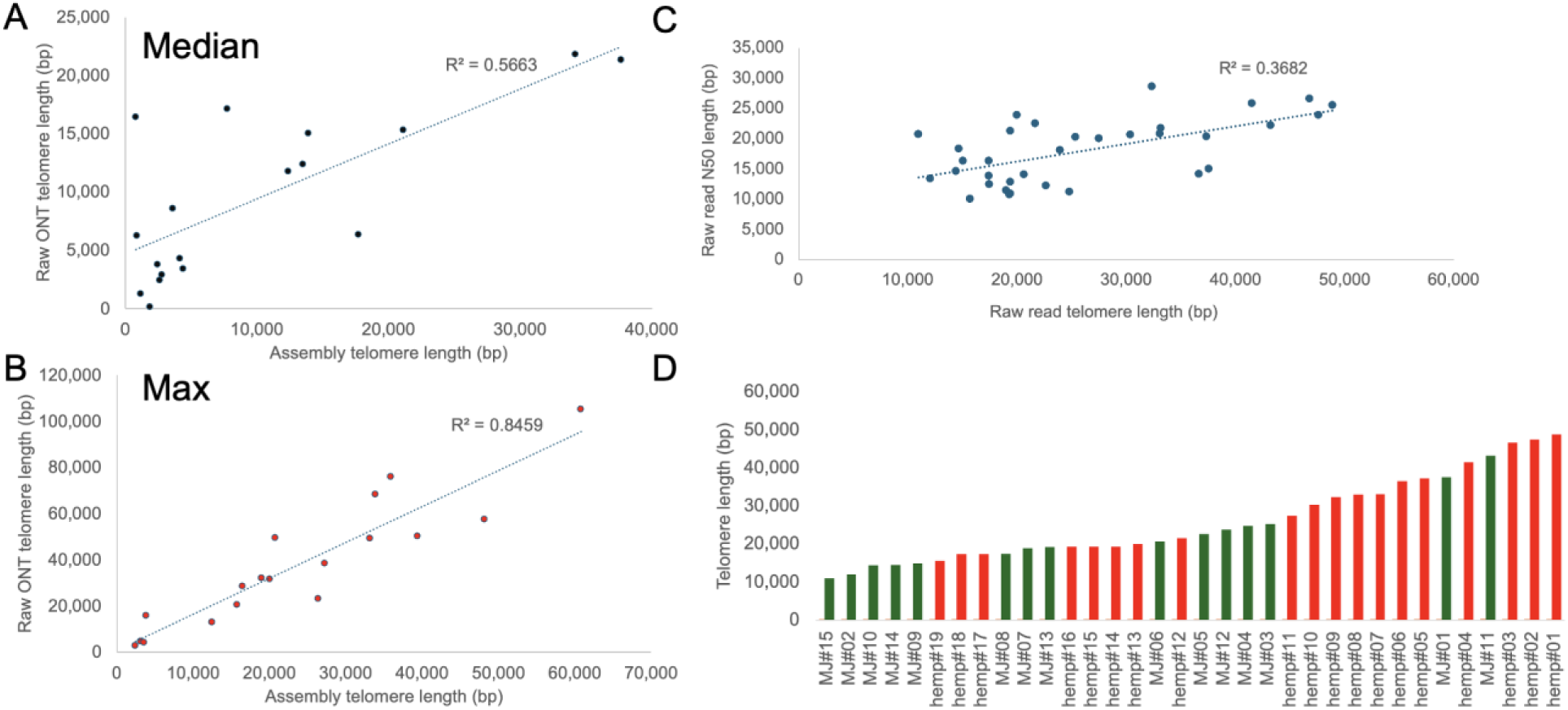
Estimating telomere length directly from raw long reads. A) Median telomere length estimated in both raw Oxford Nanopore Technologies (ONT) reads or an assembly based on the same reads. B) Maximum telomere length estimated in both raw ONT reads or an assembly based on the same reads. C) Raw read telomere length by raw read N50 length across a population of *Cannabis sativa* hemp or marijuana (MJ). D) Maximum telomere length across hemp (red) and MJ (green) lines (Supplemental Table S3).

One of the main features determining telomere estimation from raw reads was the read N50 length. Since *Cannabis sativa* had long telomeres we tested whether raw read N50 impacted the telomere length estimation. We identified a panel of both “hemp” and marijuana (MJ) lines to sample both putatively more “wild” and “domesticated” Cannabis lines respectively. We found that maximum telomere length estimates were minimally correlated with read length N50 (0.37; Figure 5C), while mean length was not correlated at all (data not shown). This suggests that read length N50 does impact maximum telomere estimation but that there are other factors such as DNA preparation and quality that also most likely play a role. However, the read length N50 was high for this entire dataset with the lowest being in 10 kb range, which is still within the same range as the estimated telomere lengths; it is possible for datasets with raw read N50 lengths below 10 kb may have a greater impact of estimating the telomere length properly. In our Cannabis panel we did note that of the hemp and MJ lines we surveyed, the hemp in general had longer telomeres and that there was a high level of variability across lines as has been noted in other species (Figure 5D).

Next we estimated the telomere length from raw plant ONT reads that were uploaded to the NCBI Short Read Archive (SRA) (Methods). We downloaded all SRA datasets that were from plants run on the ONT promethION and metadata concerning life style and type of plant was obtained online (Supplemental Table S4). We ran TeloNum on these datasets and the reads were high quality with a raw median N50 length of 24 kb and 159 telomere reads (Supplemental Table S4). We filtered out datasets with less than 100 telomere reads and also an N50 below 10 kb and plotted these species (Figure 6). We found that the median and maximum telomere prediction was largely correlated (0.76), which was consistent with our curated datasets (Figure 6A). Also, the read length N50 was not correlated with the maximum read length suggesting these estimates were high quality and not limited by the input read N50 length (Figure 6B). The telomere lengths spanned 1 kb to 60 kb and didn’t seem to follow any specific trend (Figure 6C). We found that TeloNum was a quick and effective method for predicting telomere length from long read ONT data.

**Figure 6.**
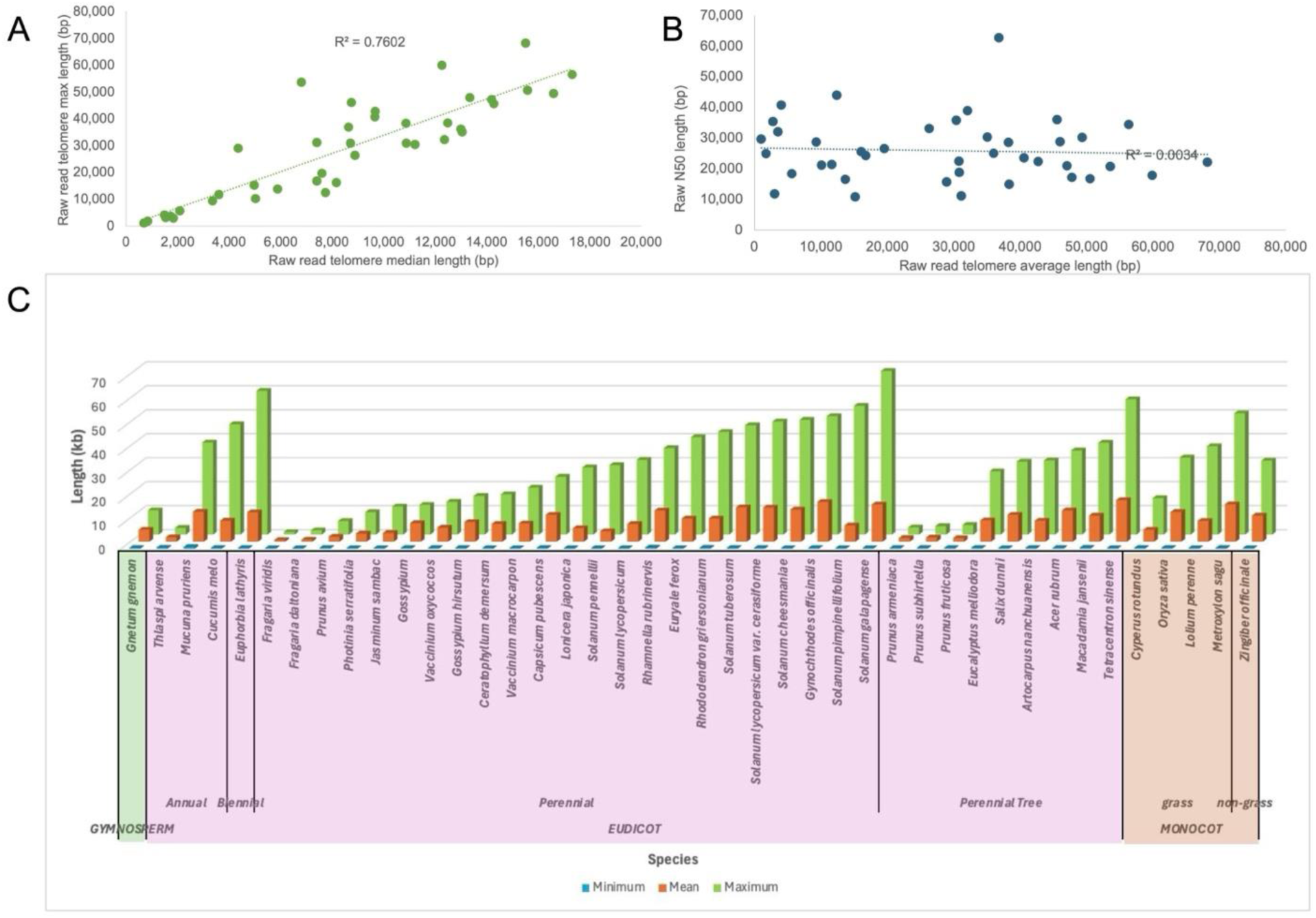
TeloNum analysis of plant telomere lengths from Oxford Nanopore Technologies (ONT) raw sequence in SRA. A) Raw ONT reads maximum versus median length based on TeloNum prediction. B) Raw read N50 length versus the maximum predicted telomere length. C) Maximum telomere from raw ONT reads sorted by maximum telomere length (Supplemental Table S4).

### Telomere length differences between wild relatives and crops

Based on the fact that we observed that hemp had longer telomeres in general compared to MJ (Supplemental Table S3), led us to hypothesize that there could be a relationship between telomere length and cultivation status: wild relatives versus crops. While we realize that hemp is also a crop it is largely an unimproved crop compared to MJ, which has undergone intense breeding historically as well as over the last 50 years. We leveraged our entire dataset including the publicly available telomere length estimations (Figure 1; Supplemental Table S1), and the new estimates with our assemblies (Figure 4,5; Supplemental Table S2) and the raw data from SRA (Figure 6; Supplemental Table S4) to look at the telomere length of wild relatives versus crops. There is a trend that crops have longer telomeres compared to their wild relatives (Figure 7).

**Figure 7.**
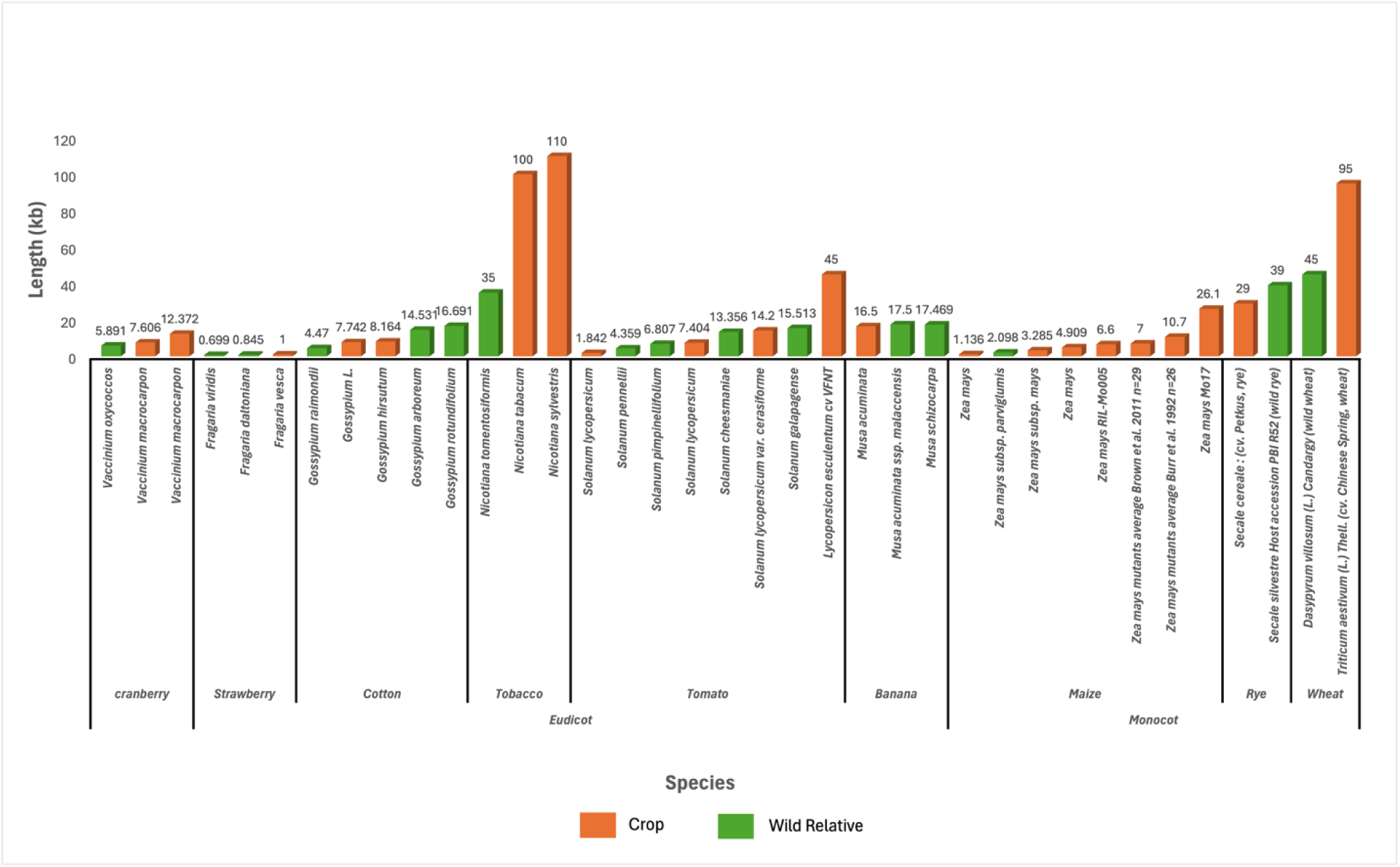
Comparison of mean telomere lengths in wild and cultivated plants. Data sourced from published telomere lengths (Supplemental Table S1), TeloNum results using raw data from SRA (Supplemental Table S2), and estimates from long read genome assemblies (Supplemental Table S4).

## Discussion

In our review of published telomere lengths in plants, patterns started to emerge amongst different lifestyle strategies. It has been shown that telomere length correlates with changes in reproductive strategy and phenotype, with the trend translating across both eudicots and monocots (Choi *et al*., 2021). Computational analysis revealed that certain arabidopsis and maize accessions, and temperate-adapted rice were found to have longer telomeres associated with faster development and earlier flowering than other cultivars (Choi *et al*., 2021). Recently Arabidopsis Col-0 was modified experimentally to possess degrees of longer and shorter telomeres than wild type; telomere length had a proportional effect on plant reproductive and anatomical traits (Campitelli *et al*., 2022). Shorter telomeres in Col-0 resulted in increased seed output at native growing conditions, while longer telomeres in Col-0 appeared to provide reproductive and fertility advantages under certain environmental stressors, with longer time until first flower (Campitelli *et al*., 2022). There are experimental differences between these studies that could be responsible for the opposite effects on flowering time (flowering data collected at different temperatures, mutants vs natural accessions); however, it’s possible that the modification of telomeres to be longer or shorter produces accession specific changes in potentially useful agronomic traits. Is there some adaptive advantage to longer or shorter telomeres that have been unknowingly selected for in the historical breeding process of crop species or specific accessions (Figure 7)? More data is needed on crops and their wild relatives before drawing any conclusions however this could be an interesting area for further study.

There are important caveats to interpreting telomere lengths generated using various methods and combinations of TRF, qPCR, Illumina, ONT, and PacBio which make comparison difficult (Supplemental Table S1). Highlighting some of the issues with these methods, (Peška *et al*., 2015) compared TRF analysis of *Cestrum elegans* telomeres, with a less common repeat motif (TTTTTTAGGG)n, using different endonucleases *Taq* I and *Sfi* I. Each endonuclease generated different size telomere fragments, 48-97kb and 48-500kb respectively, likely due to the variation in restriction enzyme binding sites across telomeric arrays. They also assessed telomere size using Illumina NGS data, with a resulting mean telomere length around 19kb, completely outside of the range estimated using TRF (Peška *et al*., 2015). This shows the significant differences and potential issues when using historic methods to estimate telomere length; it’s possible that reported telomere lengths based on older methods are inaccurate thus muddying the comparison across data types.

Recent T2T assemblies have been generated using combinations of Oxford Nanopore, PacBio HiFi, Bionano, and Illumina data with published telomere lengths for many plants: *Glycine max* Zhonghuang 13 (Zhang *et al*., 2023), *Citrullus lanatus* G42 (Deng *et al*., 2022), *Fragaria vesca* (Zhou *et al*., 2023), *Momordica charantia* L. var. *abbreviata* Ser. (Mca) (Fu *et al*., 2023), *Actinidia chinensis* HY4P and HY4A (Yue *et al*., 2023), *Zea mays* Mo17 (Chen *et al*., 2023), *Dianthus caryophyllus (Lan et al., 2024), Ficus hispida (Liao et al., 2024), Vaccinium duclouxii (Zeng et al., 2023), Vitis vinifera* Thompson Seedless (X. Wang *et al*., 2024)*, Cucumis melo* L. var. *Inodorus* (Wei *et al*., 2023)*), Mangifera indica* ‘Irwin’ (Henry et al. preprint), *Musa acuminata* Wild type (Liu *et al*., 2023) and Cavendish (Huang *et al*., 2023), *Oryza sativa* ZS97RS3 and MH63RS3 (Song *et al*., 2021), *Mycrica rubra* (Zhang *et al*., 2024) (Supplemental Table S1). However, assembled telomere sequence is often used as a means to an end in confirming a complete assembly; their details are often relegated to supplemental materials or in many cases not clearly reported at all (Belser *et al*., 2021; Bao *et al*., 2023; He *et al*., 2023; Sato *et al*., 2023; Sun *et al*., 2023; Wang *et al*., 2023; Bi *et al*., 2024; Z. Wang *et al*., 2024). NGS bioinformatic methods have been developed to detect telomeres in a T2T assembled genome of choice, for instance quarTET TeloExplorer (Lin *et al*., 2023) and TIDK v0.2.1 (Henry *et al*., 2023); The computational tools here will make it possible to broadly test the length of telomeres across a diverse array of plants, tissue types and developmental times from raw sequencing reads.

Here we validate using raw long reads directly to accurately predict telomere length as well as long read assemblies to determine chromosome variation in telomere length in *A. thaliana* and rice, which have high quality estimates across several orthologous methods. We found the longest max telomeres directly from raw reads as compared to genome assemblies (Figure 5). This is very clear in the median telomere length (Figure 5A), where almost all of the telomeres are longer from raw reads. In contrast, for the max reads only the very long telomeres are longer by raw reads (Figure 5B). These results suggest that very long telomere arrays are found in the raw data that may not get included in the assembly. This could be due to the assembly algorithms used, and that the long telomere arrays are not incorporated since we do not see this as much with the smaller telomere arrays. We recently developed a methodology to incorporate telomere sequence into an assembly that was not captured by the assembly process (Naish *et al*., 2021). We expect that as long read sequencing matures further and assembly algorithms are designed to be more repeat aware, especially for plants, assembly of all telomeres will become standard practice.

One potential benefit of estimating the telomere length from long read assemblies over other methods, including from short read technologies or raw long reads, is the association of telomeres to specific chromosomes. In the Columbia accession of *A. thaliana* the telomere length is relatively similar across the chromosomes (Avg=2,760 bp; Max=3,088; Figure 2). We found a similar situation across the chromosomes of basmati rice, which was mostly conserved between two different accessions basmati334 and Dom Sufid (Figure 3). However, only 50% (5/10) of the telomeres were assembled in *A. thaliana*, 75% (15/20) in basmati, and 36%(8/22) in banana (Figure 2,3; Supplemental Figure S2). In *A. thaliana*, the telomere is proximal to the two rDNA arrays on chromosomes 2 and 4, which makes them difficult to assemble (Naish *et al*., 2021). There is an added difficulty in assembling telomeres when there are long sub-telomeric repeats (Fajkus *et al*., 1995), although the read N50 length also plays a role in assembling these complex regions especially in a species like banana where the telomeres are on average 17 kb (Figure 5C; Supplemental Figure S2). An updated assembly of the banana genome with 17X reads greater than 75 kb enabled the identification of telomeres on all of the chromosomes with an average length of 17 kb (Belser *et al*., 2021).

While long reads and long read assemblies have the potential to broaden our understanding of telomere length across plants, there are some caveats to this approach. Long read sequencing technologies such as ONT are highly dependent on the quality of the input DNA. For annual species like *A. thaliana* where the telomeres are short (2-5 kb) we are very confident that we are getting complete telomeres. However, for species with telomeres that exceed read length (N50 length > 20 kb) like Cannabis, there is a chance that using long reads could potentially underestimate length (Figure 5B); this would be very dependent on the sequencing run and the input DNA. Our raw read pipeline, TeloNum does require some unique genome sequences, with the idea that they would ensure almost complete telomeres. It is possible that we also might miss telomeres that are not proximal to a unique sequence (e.g. sub telomeric arrays, transposable elements), or where the telomere length exceeds the read length. For these reasons, we are more confident with our longest telomeres (maximum), yet less confident when they are very short, which is likely the result of underestimated size. Another caveat is the tissue sampled for sequencing; we currently do not know if the telomere length would differ by different tissues from our study, since most of the material presented was from bulk leaf tissue.

Leveraging estimation from long reads we revealed the variability of telomere length in plants with various lifestyle habits. A notable trend was that some of the longest lived gymnosperms eg. *Welwitschia mirabilis* had substantially shorter median telomere lengths compared to perennial eudicots. *Welwitschia* in particular has an expansion of genes associated with DNA and protein integrity (Wan *et al*., 2021). In hemp we found some of the longest telomere arrays (Figure 4, 5); hemp has strong sub-telomeric repeats called CS-1 (Divashuk *et al*., 2014), which have been shown to be important for homologous chromosome pairing during meiosis in other plants (Calderón *et al*., 2014) and are remnant centromere sequences (Grassa *et al*., 2021). *Lemna*, *Wolffia* and *Wolffiella* arrays were similar to other monocots (sorghum and banana), yet were larger than other genera in the family such as *Spirodela*. Hemp/MJ and duckweed do share a common growth habit; they both grow clonally, naturally in the latter, and for crop production and consistency in the former. It is possible that organisms that predominantly are maintained clonally develop longer telomere arrays over time to protect chromosome structure. For instance, Barley telomere length dramatically increases in long term callus culture (Kilian, Stiff and Kleinhofs, 1995). However, it has been shown that in Norway Spruce telomeres are initially maintained but then shorten in tissue culture (Aronen, Virta and Varis, 2021). Care should also be taken to sample consistent tissue types for analysis; the variability of telomere length seen in duckweed could be in part due to the fact that whole plants were used in aggregate for DNA isolation. Whole duckweed fronds contain many generations of daughter fronds, meaning numerous tissue types and tissue ages were analyzed at the same time. Further studies in clonal populations and tissue culture across species are required to understand this process, and it is possible that it is species specific. The ability to rapidly estimate the length and structure of the telomere across plants using long reads and long read assemblies enables a new understanding of chromosome structure and function.

## Methods

### DNA Isolation

Plant tissue was ground by mortar and pestle into a fine powder in liquid nitrogen. Approximately 3 grams of ground sample were resuspended in 40mL of nuclei isolation buffer (NIB) composed of 15mM Tris-HCl pH 9.5, 10mM EDTA, 130mM KCl, 20mM NaCl, 8% PVP-10, 1.15mM spermine, 1.15mM spermidine, 0.10% Triton X-100, and 7.5% BME. Resuspended samples were filtered through a 100µM cell strainer, then through a 40µM cell strainer. Cell debris was pelleted at 60xg for 2 minutes. The supernatant was retained in a new tube, and centrifuged at 2,500xg for 45 minutes. The supernatant was discarded and the nuclei pellets were gently resuspended in 10mL of NIB and pelleted at 2,500xg repeatedly, until each nuclei pellet was white. The nuclei were then suspended in 0.8X low melt agarose gel plugs (BIORAD CHEF Disposable Plug Molds). After solidifying, the plugs were placed in 5mL lysis buffer, composed of 1% sodium lauryl sarcosine and 0.5M EDTA pH8. The tubes were incubated at 50°C with 200µL of 20mg/mL Proteinase K overnight. The lysis buffer and proteinase K were then removed and replaced, and a second Proteinase K treatment incubated at 50°C for two hours. The samples were then cooled to 37°C, and 50µL of 100mg/mL RNase A was added and incubated for 1 hour. The plugs were then washed by gentle rocking in 5mL of 0.5M EDTA pH 8 for a total of four, fifteen minute incubations. Then the plugs were washed by gentle rocking in 10mL of 100mM tris for a total of five, fifteen minute incubations. The buffer was removed completely and the gel plugs were placed into 1.5mL tubes. The gel plugs were incubated at 70°C until fully melted, and cooled down to 42°C for five minutes. 2µL of Beta Agarase per gel plug was added to each tube, mixed gently by flicking, and incubated for 45 minutes. The samples were then centrifuged at 16,000xg for 30 minutes, and the clear supernatant containing DNA was retained for analysis.

### Telomere identification from assembled genomes

ONT and PacBio based long read-based assemblies enable the assembly of some of the highly repetitive centromeres and telomeres sequences (VanBuren *et al*., 2015). Telomeres were identified by searching long read genomes using tandem repeat finder (TRF; v4.09) using modified settings (1 1 2 80 5 200 2000 -d -h) (Benson, 1999). Repeats with the period of seven (7) were queried for the 14 different versions of the canonical telomere base repeat: AAACCCT, AACCCTA, ACCCTAA, CCCTAAA, CCTAAAC, CTAAACC, TAAACCC, TTTAGGG, TTAGGGT, TAGGGTT, AGGGTTT, GGGTTTA, GGTTTAG, GTTTAGG. (Supplemental Table S2)

### Estimating telomere length directly from raw long ONT reads

We developed a software pipeline to estimate telomere length from raw long reads called TeloNum: https://gitlab.com/semarpetrus/telonum. Long ONT raw reads were mapped to a canonical telomere sequence fasta, and reads that were 1) larger than 10 kb; and 2) had telomere sequence in the first 1 kn or the last kb. The resulting reads were evaluated by tandem repeat finder (TRF) for telomere arrays, and telomere features were summarized along with the read length N50 (Supplemental Table S3).

### Estimating telomere length from raw reads on SRA

Long-read PromethION datasets were identified on the sequencing read archive (SRA) at the National Center for Biotechnology Information (NCBI) on 11/02/2021. Sequencing datasets were merged if partial fastq files were uploaded, and datasets with less than 100 reads were removed from the analysis. The TeloNum results were further filtered for datasets that had more than 100 telomere reads, and a raw N50 read length greater than 10 kb (Supplemental Table S4).

### Monochrome multiplex quantitative PCR (mmqPCR)

Monochrome multiplex quantitative PCR was performed based on the methods described in (Vaquero-Sedas and Vega-Palas, 2014). To amplify telomere sequences, the following primers were used from (Vaquero-Sedas and Vega-Palas, 2014): “TelA” 5’-CCCCGGTTTTGGGTTTTGGGTTTTGGGTTTTGGGT-3’ Tm=79°C and “TelB” 5’-GGGGCCCTAATCCCTAATCCCTAATCCCTAATCCCT-3’ Tm=77°C. We selected two single copy genes (SCG) GIGANTEA and LFY (for arabidopsis CYP5 was also included). For each plant, primers were designed in unique regions of the single copy genes to avoid off target amplification. GC tails were added to the 5’ ends of the SCG primers to raise the Tm to ∼95°C for multiplexing. SCG primers are included in the supplemental materials (Supplemental Table S5). For each of the following primer combinations, “SCG only”, “telomere only”, and “SCG+telomere”; standard curves were made for each sample using a serial dilution of gDNA from 100 to 0.01ng. Primers were diluted to 10mM in 10mM Tris pH 8. qPCR reactions were set up as follows: 0.75 uL water, 2.5 uL 10X sybr green 1 (Life Technologies), 2.5 uL ImmoBuffer™ (Bioline) 103, 1.25 uL MgCl2 50mM, 2 uL dNTPs mix 2.5 mM, 1.5 uL each 10uM primer, 0.125 uL Immolase™ DNA polymerase (Bioline), 10 uL DNA for a total of 10ng input, and if only one set of primers were used added 3 uL10mM Tris pH 8 to account for volume. All standards and samples were run in triplicate. PCR cycling conditions were as follows: Stage 1) 15 minutes at 95°C; Stage 2) Two cycles of 15 seconds at 94°C and 15 seconds at 49°C; Stage 3) 32 cycles of: 15 seconds at 94°C, 10 seconds at 62°C, 15 seconds at 74°C with signal acquisition (telomeric Ct values captured), 10 seconds at 84°C, 15 seconds at 85°C with signal acquisition (SCG Ct values captured). Samples were run on the Bio-Rad CFX96 Touch Deep Well Real-Time PCR Detection System and results were exported as .csv files from CFX Maestro™ Software. To estimate the telomere length for each species, we used the following equation: [(2^(SCG Ct - telomere Ct))/ploidy].

## Author contributions

TPM conceived the study. KC and BWA sequenced genomes. SP and NTH assembled genomes. SP developed TeloNum. TPM analyzed telomeres. KC carried out validation. TPM and KC wrote the manuscript with all authors providing edits.

## Acknowledgements

We thank Michael lab members for their comments and insight, and Tom Kursar and Lissy Coley for Inga leaf tissue.

## References

Abdulkina, L.R. et al. (2019) ‘Components of the ribosome biogenesis pathway underlie establishment of telomere length set point in Arabidopsis’, Nature communications, 10(1), p. 5479.

Abramson, B.W. et al. (2021) ‘The genome and preliminary single-nuclei transcriptome of Lemna minuta reveals mechanisms of invasiveness’, Plant physiology [Preprint]. Available at: 10.1093/plphys/kiab564.

Alonge, M. et al. (2019) ‘RaGOO: fast and accurate reference-guided scaffolding of draft genomes’, Genome biology, 20(1), p. 224.

Alonge, M. et al. (2022) ‘Automated assembly scaffolding using RagTag elevates a new tomato system for high-throughput genome editing’, Genome biology, 23(1), p. 258.

Aronen, T. and Ryynänen, L. (2012) ‘Variation in telomeric repeats of Scots pine (Pinus sylvestris L.)’, Tree genetics & genomes, 8(2), pp. 267–275.

Aronen, T., Virta, S. and Varis, S. (2021) ‘Telomere Length in Norway Spruce during Somatic Embryogenesis and Cryopreservation’, Plants, 10(2). Available at: 10.3390/plants10020416.

Aubert, G., Hills, M. and Lansdorp, P.M. (2012) ‘Telomere length measurement-caveats and a critical assessment of the available technologies and tools’, Mutation research, 730(1-2), pp. 59–67.

Baggs, E.L. et al. (2022) ‘Characterization of defense responses against bacterial pathogens in duckweeds lacking EDS1’, The New phytologist, 236(5), pp. 1838–1855.

Bao, Y. et al. (2023) ‘A gap-free and haplotype-resolved lemon genome provides insights into flavor synthesis and huanglongbing (HLB) tolerance’, Horticulture research, 10(4), p. uhad020.

Barcenilla, B.B. et al. (2023) ‘Arabidopsis telomerase takes off by uncoupling enzyme activity from telomere length maintenance in space’, Nature communications, 14(1), p. 7854.

Belser, C. et al. (2018) ‘Chromosome-scale assemblies of plant genomes using nanopore long reads and optical maps’, Nature plants, 4(11), pp. 879–887.

Belser, C. et al. (2021) ‘Telomere-to-telomere gapless chromosomes of banana using nanopore sequencing’, bioRxiv. Available at: 10.1101/2021.04.16.440017.

Benson, G. (1999) ‘Tandem repeats finder: a program to analyze DNA sequences’, Nucleic acids research, 27(2), pp. 573–580.

Bi, G. et al. (2024) ‘Near telomere-to-telomere genome of the model plant Physcomitrium patens’, Nature Plants, 10(2), pp. 327–343.

Brown, A.N. et al. (2011) ‘QTL Mapping and Candidate Gene Analysis of Telomere Length Control Factors in Maize (Zea mays L.)’, G3: *Genes, Genomes, Genetics*, 1(6), pp. 437–450.

Burr, B. et al. (1992) ‘Pinning down loose ends: mapping telomeres and factors affecting their length’, The Plant cell, 4(8), pp. 953–960.

Calderón, M. del C., et al. (2014) ‘The subtelomeric region is important for chromosome recognition and pairing during meiosis’, Scientific reports, 4, p. 6488.

Campitelli, B.E. et al. (2022) ‘Plasticity, pleiotropy and fitness trade-offs in Arabidopsis genotypes with different telomere lengths’, The New phytologist, 233(4), pp. 1939–1952.

Cawthon, R.M. (2009) ‘Telomere length measurement by a novel monochrome multiplex quantitative PCR method’, Nucleic acids research, 37(3), p. e21.

Chen, J. et al. (2023) ‘A complete telomere-to-telomere assembly of the maize genome’, Nature genetics, 55(7), pp. 1221–1231.

Choi, J.Y. et al. (2020) ‘Nanopore sequencing-based genome assembly and evolutionary genomics of circum-basmati rice’, Genome biology, 21(1), p. 21.

Choi, J.Y. et al. (2021) ‘Natural variation in plant telomere length is associated with flowering time’, The Plant cell [Preprint]. Available at: 10.1093/plcell/koab022.

Cowan, C.R., Carlton, P.M. and Cande, W.Z. (2001) ‘The polar arrangement of telomeres in interphase and meiosis. Rabl organization and the bouquet’, Plant physiology, 125(2), pp. 532– 538.

Deng, Y. et al. (2022) ‘A telomere-to-telomere gap-free reference genome of watermelon and its mutation library provide important resources for gene discovery and breeding’, Molecular plant, 15(8), pp. 1268–1284.

Deschamps, S. et al. (2018) ‘A chromosome-scale assembly of the sorghum genome using nanopore sequencing and optical mapping’, Nature communications, 9(1), p. 4844.

Divashuk, M.G. et al. (2014) ‘Molecular Cytogenetic Characterization of the Dioecious Cannabis sativa with an XY Chromosome Sex Determination System’, PLoS ONE, p. e85118. Available at: 10.1371/journal.pone.0085118.

Fajkus, J. et al. (1995) ‘Organization of telomeric and subtelomeric chromatin in the higher plant Nicotiana tabacum’, Molecular & general genetics: MGG, 247(5), pp. 633–638.

Fajkus, J., Sýkorová, E. and Leitch, A.R. (2005) ‘Techniques in plant telomere biology’, BioTechniques, 38(2), pp. 233–243.

Fajkus, P. et al. (2016) ‘Allium telomeres unmasked: the unusual telomeric sequence (CTCGGTTATGGG)n is synthesized by telomerase’, The Plant journal: for cell and molecular biology, 85(3), pp. 337–347.

Flanary, B.E. and Kletetschka, G. (2006) ‘Analysis of telomere length and telomerase activity in tree species of various lifespans, and with age in the bristlecone pine Pinus longaeva’, Rejuvenation research, 9(1), pp. 61–63.

Fojtová, M. et al. (2015) ‘Terminal restriction fragments (TRF) method to analyze telomere lengths’, Bio-protocol, 5(23), pp. e1671–e1671.

Fu, A. et al. (2023) ‘Telomere-to-telomere genome assembly of bitter melon (Momordica charantia L. var. abbreviata Ser.) reveals fruit development, composition and ripening genetic characteristics’, Horticulture research, 10(1), p. uhac228.

Fulcher, N. et al. (2015) ‘Genetic architecture of natural variation of telomere length in Arabidopsis thaliana’, Genetics, 199(2), pp. 625–635.

Ganal, M.W., Lapitan, N.L. and Tanksley, S.D. (1991) ‘Macrostructure of the tomato telomeres’, The Plant cell, 3(1), pp. 87–94.

Gomes, N.M.V. et al. (2011) ‘Comparative biology of mammalian telomeres: hypotheses on ancestral states and the roles of telomeres in longevity determination’, Aging cell, 10(5), pp. 761–768.

Grassa, C.J. et al. (2021) ‘A new Cannabis genome assembly associates elevated cannabidiol (CBD) with hemp introgressed into marijuana’, The New phytologist, 230(4), pp. 1665–1679.

Harkess, A. et al. (2020) ‘A new Spirodela polyrhiza genome and proteome reveal a conserved chromosomal structure with high abundances of proteins favoring energy production’. Available at: 10.1101/2020.01.23.909457.

Harkess, A. et al. (2021) ‘Improved Spirodela polyrhiza genome and proteomic analyses reveal a conserved chromosomal structure with high abundance of chloroplastic proteins favoring energy production’, Journal of experimental botany, 72(7), pp. 2491–2500.

Henry, R. et al. (2023) ‘A telomere-to-telomere genome of mango exclusively from long-read sequence data’, Research Square. Available at: 10.21203/rs.3.rs-3588192/v1.

He, S. et al. (2023) ‘A telomere-to-telomere reference genome provides genetic insight into the pentacyclic triterpenoid biosynthesis in Chaenomeles speciosa’, Horticulture research, 10(10), p. uhad183.

Hoang, P.N.T. et al. (2018) ‘Generating a high-confidence reference genome map of the Greater Duckweed by integration of cytogenomic, optical mapping, and Oxford Nanopore technologies’, The Plant journal: for cell and molecular biology, 96(3), pp. 670–684.

Huang, H.-R. et al. (2023) ‘Telomere-to-telomere haplotype-resolved reference genome reveals subgenome divergence and disease resistance in triploid Cavendish banana’, Horticulture research, 10(9), p. uhad153.

Kilian, A., Stiff, C. and Kleinhofs, A. (1995) ‘Barley telomeres shorten during differentiation but grow in callus culture’, Proceedings of the National Academy of Sciences of the United States of America, 92(21), pp. 9555–9559.

Kumawat, S. and Choi, J.Y. (2023) ‘No end in sight: Mysteries of the telomeric variation in plants’, American journal of botany, 110(11), p. e16244.

Lan, L. et al. (2024) ‘The haplotype-resolved telomere-to-telomere carnation (Dianthus caryophyllus) genome reveals the correlation between genome architecture and gene expression’, Horticulture research, 11(1), p. uhad244.

Liao, Z. et al. (2024) ‘A telomere-to-telomere reference genome of ficus (Ficus hispida) provides new insights into sex determination’, Horticulture research, 11(1), p. uhad257.

Lin, Y. et al. (2023) ‘quarTeT: a telomere-to-telomere toolkit for gap-free genome assembly and centromeric repeat identification’, Horticulture research, 10(8), p. uhad127.

Liu, D. et al. (2007) ‘Comparative analysis of telomeric restriction fragment lengths in different tissues of Ginkgo biloba trees of different age’, Journal of plant research, 120(4), pp. 523–528.

Liu, X. et al. (2023) ‘The phased telomere-to-telomere reference genome of Musa acuminata, a main contributor to banana cultivars’, Scientific data, 10(1), p. 631.

Maheshwari, S. et al. (2017) ‘Centromere location in Arabidopsis is unaltered by extreme divergence in CENH3 protein sequence’, Genome research, 27(3), pp. 471–478.

Michael, T.P. et al. (2018) ‘High contiguity Arabidopsis thaliana genome assembly with a single nanopore flow cell’, Nature communications, 9(1), p. 541.

Michael, T.P. et al. (2020) ‘Genome and time-of-day transcriptome of Wolffia australiana link morphological minimization with gene loss and less growth control’, Genome research [Preprint]. Available at: 10.1101/gr.266429.120.

Mizuno, H. et al. (2006) ‘Sequencing and characterization of telomere and subtelomere regions on rice chromosomes 1S, 2S, 2L, 6L, 7S, 7L and 8S’, The Plant journal: for cell and molecular biology, 46(2), pp. 206–217.

Mizuno, H., Matsumoto, T. and Wu, J. (2018) ‘Composition and Structure of Rice Centromeres and Telomeres’, in T. Sasaki and M. Ashikari (eds) Rice Genomics, Genetics and Breeding. Singapore: Springer Singapore, pp. 37–52.

Mu, Y. et al. (2014) ‘Sex-and season-dependent differences in telomere length and telomerase activity in the leaves of ash and willow’, SpringerPlus, 3, p. 163.

Naish, M. et al. (2021) ‘The genetic and epigenetic landscape of the Arabidopsis centromeres’, bioRxiv. Available at: 10.1101/2021.05.30.446350.

Osnato, M. (2021) ‘Searching for the link between telomere length and life history traits in plants’, The Plant cell, 33(4), pp. 1087–1088.

Paez, M.Z., Holliday, J. and Hamilton, J. (2023) ‘Leveraging whole genome sequencing to estimate telomere length in plants’, Authorea Preprints [Preprint]. Available at: 10.22541/au.168569940.01641234/v1.

Peška, V. et al. (2015) ‘Characterisation of an unusual telomere motif (TTTTTTAGGG)n in the plant Cestrum elegans (Solanaceae), a species with a large genome’, The Plant journal: for cell and molecular biology, 82(4), pp. 644–654.

Peska, V. and Garcia, S. (2020) ‘Origin, Diversity, and Evolution of Telomere Sequences in Plants’, Frontiers in plant science, 11, p. 117.

Peska, V., Sýkorová, E. and Fajkus, J. (2008) ‘Two faces of Solanaceae telomeres: a comparison between Nicotiana and Cestrum telomeres and telomere-binding proteins’, Cytogenetic and genome research, 122(3-4), pp. 380–387.

Procházková Schrumpfová, P., Fojtová, M. and Fajkus, J. (2019) ‘Telomeres in Plants and Humans: Not So Different, Not So Similar’, Cells, 8(1), p. 58.

Richards, E.J. and Ausubel, F.M. (1988) ‘Isolation of a higher eukaryotic telomere from Arabidopsis thaliana’, Cell, 53(1), pp. 127–136.

Risques, R.A. and Promislow, D.E.L. (2018) ‘All’s well that ends well: why large species have short telomeres’, Philosophical transactions of the Royal Society of London. Series B, Biological sciences, 373(1741). Available at: 10.1098/rstb.2016.0448.

Sato, M.P. et al. (2023) ‘Telomere-to-telomere genome assembly of an allotetraploid pernicious weed, Echinochloa phyllopogon’, DNA research: an international journal for rapid publication of reports on genes and genomes, 30(5). Available at: 10.1093/dnares/dsad023.

Shakirov, E.V. and Shippen, D.E. (2004) ‘Length regulation and dynamics of individual telomere tracts in wild-type Arabidopsis’, The Plant cell, 16(8), pp. 1959–1967.

Shan-An, H., Gu, Y. and Zi-Jie, P. (1997) ‘Resources and Prospects of Ginkgo biloba in China’, in T. Hori et al. (eds) Ginkgo Biloba A Global Treasure: From Biology to Medicine. Tokyo: Springer Japan, pp. 373–383.

Song, J.-M. et al. (2021) ‘Two gap-free reference genomes and a global view of the centromere architecture in rice’, Molecular plant, 14(10), pp. 1757–1767.

Soreng:, R.J. and Peterson, P.M. (no date) Tropicos. Available at: http://legacy.tropicos.org/Project/CNWG (Accessed: 13 March 2024).

Stindl, R. (2014) ‘The telomeric sync model of speciation: species-wide telomere erosion triggers cycles of transposon-mediated genomic rearrangements, which underlie the saltatory appearance of nonadaptive characters’, Die Naturwissenschaften, 101(3), pp. 163–186.

Sun, M. et al. (2023) ‘Telomere-to-telomere pear (Pyrus pyrifolia) reference genome reveals segmental and whole genome duplication driving genome evolution’, Horticulture research, 10(11), p. uhad201.

Sykorova, E. et al. (2003) ‘The absence of Arabidopsis-type telomeres in Cestrum and closely related genera Vestia and Sessea (Solanaceae): first evidence from eudicots’, The Plant journal: for cell and molecular biology, 34(3), pp. 283–291.

Tan, K.-T. et al. (2022) ‘Identifying and correcting repeat-calling errors in nanopore sequencing of telomeres’, Genome biology, 23(1), p. 180.

Tran, T.D. et al. (2015) ‘Centromere and telomere sequence alterations reflect the rapid genome evolution within the carnivorous plant genus Genlisea’, The Plant journal: for cell and molecular biology, 84(6), pp. 1087–1099.

VanBuren, R. et al. (2015) ‘Single-molecule sequencing of the desiccation-tolerant grass Oropetium thomaeum’, Nature, 527(7579), pp. 508–511.

Vaquero-Sedas, M.I. and Vega-Palas, M.A. (2014) ‘Determination of Arabidopsis thaliana telomere length by PCR’, Scientific reports, 4, p. 5540.

Vershinin, A.V. and Heslop-Harrison, J.S. (1998) ‘Comparative analysis of the nucleosomal structure of rye, wheat and their relatives’, Plant molecular biology, 36(1), pp. 149–161.

Vitales, D. et al. (2017) ‘Third release of the plant rDNA database with updated content and information on telomere composition and sequenced plant genomes’, Plant systematics and evolution = Entwicklungsgeschichte und Systematik der Pflanzen, 303(8), pp. 1115–1121.

Wang, W. et al. (2020) ‘The draft nuclear genome assembly of Eucalyptus pauciflora: a pipeline for comparing de novo assemblies’, GigaScience, 9(1). Available at: 10.1093/gigascience/giz160.

Wang, X. et al. (2024) ‘Telomere-to-telomere and gap-free genome assembly of a susceptible grapevine species (Thompson Seedless) to facilitate grape functional genomics’, Horticulture research, 11(1), p. uhad260.

Wang, Y.-H. et al. (2023) ‘Telomere-to-telomere carrot (Daucus carota) genome assembly reveals carotenoid characteristics’, Horticulture research, 10(7), p. uhad103.

Wang, Z. et al. (2024) ‘Insights into the Superrosids phylogeny and flavonoid synthesis from the telomere-to-telomere gap-free genome assembly of Penthorum chinense Pursh’, Horticulture Research, 11(2). Available at: 10.1093/hr/uhad274.

Wan, T. et al. (2021) ‘The Welwitschia genome reveals a unique biology underpinning extreme longevity in deserts’, Nature communications, 12(1), p. 4247.

Wei, M. et al. (2023) ‘Telomere-to-telomere genome assembly of melon (Cucumis melo L. var. inodorus) provides a high-quality reference for meta-QTL analysis of important traits’, Horticulture research, 10(10), p. uhad189.

Weiss, H. and Scherthan, H. (2002) ‘Aloe spp.--plants with vertebrate-like telomeric sequences’, Chromosome research: an international journal on the molecular, supramolecular and evolutionary aspects of chromosome biology, 10(2), pp. 155–164.

Whittemore, K. et al. (2019) ‘Telomere shortening rate predicts species life span’, Proceedings of the National Academy of Sciences of the United States of America, 116(30), pp. 15122– 15127.

Yue, J. et al. (2023) ‘Telomere-to-telomere and gap-free reference genome assembly of the kiwifruit Actinidia chinensis’, Horticulture research, 10(2), p. uhac264.

Zeng, T. et al. (2023) ‘The telomere-to-telomere gap-free reference genome of wild blueberry (Vaccinium duclouxii) provides its high soluble sugar and anthocyanin accumulation’, Horticulture research, 10(11), p. uhad209.

Zhang, A. et al. (2023) ‘A telomere-to-telomere genome assembly of Zhonghuang 13, a widely-grown soybean variety from the original center of Glycine max’, The Crop Journal [Preprint]. Available at: 10.1016/j.cj.2023.10.003.

Zhang, S. et al. (2024) ‘T2T reference genome assembly and genome-wide association study reveal the genetic basis of Chinese bayberry fruit quality’, Horticulture research, 11(3), p. uhae033.

Zhou, Y. et al. (2023) ‘The telomere-to-telomere genome of Fragaria vesca reveals the genomic evolution of Fragaria and the origin of cultivated octoploid strawberry’, Horticulture research, 10(4), p. uhad027.

